# An integrated analysis tool reveals intrinsic biases in gene set enrichment

**DOI:** 10.1101/2021.07.12.452009

**Authors:** Nishant Thakur, Nathalie Pujol, Jacques van Helden, Robert H. Waterston, LaDeana W. Hillier, Laurent Tichit, Jonathan J. Ewbank

**Affiliations:** Aix Marseille Univ, CNRS, INSERM, CIML, CIML, Turing Centre for Living Systems, Marseille, France; Technological Advances for Genomics and Clinics (TAGC), INSERM, U1090, Aix Marseille Université, Campus de Luminy, 13288 Marseille, France; Department of Genome Sciences, School of Medicine, University of Washington, Seattle, Washington 98195, USA; Institut de Mathématiques de Marseille, Aix Marseille Université, I2M Centrale Marseille, CNRS UMR 7373, 13453 Marseille, France

## Abstract

Generating meaningful interpretations of gene lists remains a challenge for all large-scale studies. Many approaches exist, often based on evaluating gene enrichment among pre-determined gene classes. Here, we conceived and implemented yet another analysis tool (YAAT), specifically for data from the widely-used model organism *C. elegans*. YAAT extends standard enrichment analyses, using a combination of co-expression data and profiles of phylogenetic conservation, to identify groups of functionally-related genes. It additionally allows class clustering, providing inference of functional links between groups of genes. We give examples of the utility of YAAT for uncovering unsuspected links between genes and show how the approach can be used to prioritise genes for in-depth study. Our analyses revealed several limitations to the meaningful interpretation of gene lists, specifically related to data sources and the “universe” of gene lists used. We hope that YAAT will represent a model for integrated analysis that could be useful for large-scale exploration of biological function in other species.

## INTRODUCTION

In recent years, there has been a boom in genomic, transcriptomic and epigenomic studies, largely fuelled by advances in sequencing technologies and the attendant reduction in costs. They often result in the production of long lists of candidate genes. Various publicly-available resources classify genes on the basis of their structure, interactions or function. A common way to interpret a new gene list is to exploit this available knowledge, looking for enrichment of genes in defined classes. Despite inherent limitations and the tendency for results to be moulded to fit “a sensible biological narrative” (Pavlidis, et al., 2012), particularly in hypothesis-driven research (Yanai and Lercher, 2020), this technique of gene functional enrichment has been used for more than a decade (Huang da, et al., 2009) and there are currently dozens of available tool, many listed at http://omictools.com/.

There are, however, concerns about these tools, even for the most highly cited ones. A major problem with many of them is that they are not regularly updated. Among the 21 most popular tools in 2016, 12 had not been updated since 2011; 84% of citations to gene enrichment tools refer to those that are outdated (Wadi, et al., 2016). For example, the data in DAVID was not updated for 7 years, between version 6.7, made public in 2009, and version 6.8, released in 2016, and according to the DAVID website has not been updated since. Despite this, the tool is still being cited several thousand times each year. This is a problem for 2 reasons. Firstly, gene predictions change. For the model organism *Caenorhabditis elegans*, between the WS220 release of the database Wormbase in 2009 and WS255 (released in 2016), for instance, more than 1700 new genes were defined, and the predicted structure of hundreds of existing genes modified. If tools do not update their source data, an increasing number of genes may be either incorrectly associated with an annotation, or simply absent. Secondly, these tools lack up-to-date information about gene function. Recently, gene ontology (GO) annotations have increased on average by 12.5% every year (Wadi, et al., 2016). There is a similar trend for other annotations, like those from Reactome (Fabregat, et al., 2016) and KEGG (Kanehisa, et al., 2010). Thus, for example, fully 80% of functional annotations available in mid-2016 were absent from DAVID 6.7. This is clearly a severe limitation, and can substantially bias analyses of gene lists (Wadi, et al., 2016).

Some tools that perform gene enrichment analyses are regularly updated. One example is g:Profiler (Reimand, et al., 2016). Despite its power, in common with similar tools, g:Profiler captures annotations for multiple species from centralized databases, rather than leveraging the annotations available in species-specific databases. The GO data for *C. elegans*, for example, is derived from GO Consortium’s compendium, so that it almost always lags behind Wormbase releases. Further, Wormbase contains more-or-less detailed descriptions of the phenotypes associated with mutation or RNAi-knockdown for ~40% of genes. Although much of this information in time reaches generic databases, there is always a delay, so at a given moment, for some annotations, Wormbase is the sole source. Wormbase also harbours more than 1600 transcriptome expression datasets that are manually curated to a standard machine-readable format, allowing automated retrieval and data integration. This represents a powerful resource for functional enrichment analysis that is more readily exploited than the raw data available through the main transcriptome databases such as the Gene Expression Omnibus (GEO).

Thus while generic databases tools have a broad appeal, there is a clear demand for model organism databases (MODs) (Oliver, et al., 2016) and tools that are species-specific. WormExp is one of many *C. elegans*-specific bioinformatic tools. It overcomes some of the limitations of the more generic ones, but is deliberately limited in its scope of data, principally from transcriptome studies (Yang, et al., 2016). Others include a database of time-resolved expression data (Grun, et al., 2014; Stoeckius, et al., 2014), catalogues of potential transcription factor binding sites, established on the basis of the DNA conservation (cisRED (Sleumer, et al., 2009)) or ChiPseq experiments (motif-disc (Araya, et al., 2014)), tools such as Worm-Cat, GExplore and WormMine for large-scale data mining related to gene or protein function (Ding, et al., 2018; Harris, et al., 2014; Hutter, et al., 2009), Tissue Enrichment Analysis among gene sets (Angeles-Albores, et al., 2016), WormNet that can generate new members for a pathway, infer functions from network neighbours, or predict “hub” genes, on the basis of annotations and expression data (Cho, et al., 2014) and GeneModules to identify active “modules” in gene expression data (Cary, et al., 2020). On the other hand, to the best of our knowledge, there is no single tool for enrichment analysis that exploits the diverse sets of data available for *C. elegans*. As reported here, we created a tool YAAT (for “yet another analysis tool”) that performs enrichment analysis using a very extensive dataset (> 10000 classes) gathered from multiple sources. To increase its utility, we supplemented it with several complementary analytical methods, enriched class clustering, evaluation of conservation, phylogenetic and co-expression profiling, as well as tools for visual exploration of links between classes.

In the course of benchmarking to explore the robustness of test results, we revealed biases that have the potential to skew the results of broad-category enrichment analyses. Our results show how caution is needed when interpreting any result from gene enrichment analyses in *C. elegans*. These concerns are likely to be true across species.

## SYSTEM AND METHODS

### Data collection

To constitute the database for functional enrichment analysis, we expanded our previous collection of expression and phenotypic data from diverse resources (Zugasti, et al., 2016). Firstly, for expression data we extracted ~1600 Serial Pattern of Expression Levels Locator (SPELL; (Hibbs, et al., 2007)) expression clusters, curated from several hundred articles, from the Wormbase FTP site https://ftp://caltech.wormbase.org/pub/wormbase/spell_download/tables/. We supplemented this data with non-redundant datasets from WormExp (Yang, et al., 2016). As neither Wormbase expression clusters nor WormExp are comprehensive, we manually curated a further 188 datasets from 71 articles, building on our previously described in-house set derived from 69 articles (Engelmann, et al., 2011). Most of the phenotypic data was extracted from Wormbase. Thus 1907 phenotypic classes, associated with a total of 8077 genes, were downloaded from Wormbase release WS255 (https://ftp://ftp.wormbase.org/pub/wormbase/releases/WS255/ONTOLOGY/phenotype_association.WS255.wb). Further, 101 phenotypic classes were extracted from DRSC FlyRNAi database (http://www.flyrnai.org/RNAi_all_hits.txt) and the correspondence between fly genes and their nematode homologues established as previously described (Zugasti, et al., 2016). We also automatically extracted the list of genes associated with each of the 132 *C. elegans* pathways present in KEGG. Finally, we added regulatory information into YAAT by including 202 datasets for the putative gene targets for 94 transcription factors from the last modENCODE release (Araya, et al., 2014). In addition, we pulled into YAAT the gene associations for 5668 GO terms.

### Maintaining data up to date

The database for functional enrichment analysis includes categories of “static” data from sources that are not updated, such as from the DRSC FlyRNAi database, modENCODE and any transcriptome data referenced to gene not a specific sequence, and “dynamic” data, for example RNAi phenotypes from Wormbase that are referenced to specific genomic coordinates. The static data was kept up to date by tracking changes in gene structure prediction for each set, using an R script based on the algorithms underlying Wormbase Converter (Engelmann, et al., 2011). The different sources of dynamic data have their particular release schedules. Our aim was to match YAAT updates to the Wormbase release cycle, with an interval of 5 releases (http://www.wormbase.org/about/release_schedule). We were successful in completing one update cycle and therefore data were generated for WS255 and WS260. The latter is a very extensive resource of 10395 datasets.

### Enrichment analysis

For functional enrichment statistics, we used the hypergeometric distribution using the R “stats” package. The p-value is calculated as phyper(q = x-1, m = m, n = n, k = k, lower.tail = FALSE) Where k is the number of genes in the user’s query.

N is total number of genes annotated in the catalogue of reference classes (e.g. 20129 genes for the analyses of data from WS255 reported here). m is the number of genes annotated in the functional class of interest. x is number of genes in the user list that are annotated in the functional class of interest. n= N − m For multiple comparisons, p-values are corrected with the p.adjust function in the R “stats” package, with various types of correctional statistics: “holm” (Holm, 1979), “hochberg” (Hochberg, 1988), “hommel” (Hommel, 1988), “BH” or its alias “fdr” and “BY” (Benjamini and Yekutieli, 2001).

### Clustering of enriched classes

Having defined the enriched classes for the user’s list of genes, a binary matrix of enriched classes and the overlapping genes in each class is generated, where rows represent genes found in at least one enriched class and columns represents enriched classes. Genes common to the user list and the enriched class are set to be “1”, otherwise genes are assigned the value “0”. The distance between all the genes and all the enriched classes are calculated and based on the distances, hierarchical clustering is performed using “hclust” function in R.

### Constructing a network of enriched classes

Johnson2 similarity scores (sim = (a/(a+b))+(a/(a+c)); a = number of shared genes, b = number of genes only found in one class, c = number of genes only found in the other class) were calculated between each pair of enriched classes. Nodes represent the enriched functional classes and edges represent the similarity between different classes. The user sets a threshold for the maximum distance between nodes, thereby defining the number of edges. Nodes representing functional classes found in the same hierarchical cluster are given the same colour. This representation complements the heatmap display that suffers from the usual drawback of occasional incorrect clustering seen with agglomerative methods.

### Phylogenetic profiles

For phylogenetic profile analysis, using WS255 we made a reference set of 101 proteomes from 72 free-living nematode and 29 parasitic species (nematodes and platyhelminths, collectively referred to as helminths), based on the complete set of proteins predicted from genome sequences, extracted from Wormbase. With the inclusion of more species in WS260, this was subsequently expanded to 127 helminth proteomes when the YAAT database was updated. A second reference set was constructed from the proteomes of 113 species (vertebrates, invertebrates, fungi, plants protists and prokaryotes). For this, we downloaded from Ensembl/Ensemblgenomes the sequences for the proteins of 66 reference proteomes defined by the Quest For Orthologs consortium (http://questfororthologs.org/; original data release 2016/05/03 2016_04, based on UniProt Release 2016_04, Ensembl release 84 and Ensembl Genome release 31). The complete predicted protein sets for a further 57 metazoan species, chosen for their phylogenetic diversity, but including all 5 available *Caenorhabditis* species, were downloaded from Ensembl (http://metazoa.ensembl.org; release 32). Each *C. elegans* protein was compared to the complete set of proteins in each of the helminths or diverse proteomes (including *C. elegans*) using BLASTP. When multiple isoforms existed for a single protein, the longest one was selected. For each *C. elegans* query protein, only the best hit in each species was considered. Based on this analysis, for each of the 2 datasets, we generated a “BinMat” matrix (n proteins × m species), where for each entry C_ij_, if there was a BLASTP hit in species “j” for *C. elegans* protein “i” with a P-value less than 10^−5^, then C_ij_ was set to “1”, and otherwise to “0”. Users provide lists of proteins or genes. In the latter case, for each gene, the corresponding protein (the longest isoform, if applicable) is identified. Starting with this list of “N” genes, the tool will extract a “QuBinMat” matrix (N proteins × m species) from “BinMat”. Next, the distances between the “N” proteins within the QuBinMat matrix are calculated and based on these distances, clustering is performed with the “hclust” clustering algorithm in R.

To calculate an average conservation score across the N species for a given list of n genes, first, for each gene, the number of species predicted to contain an ortholog, as defined above, was determined (o_1_, o_2_,‥o_n_). Next, the sum of the number of orthologues was divided by the maximum possible score (∑_1-n_ o /Nn).

### Co-expression dataset

A coherent set of the results from 240 independent RNAseq experiments performed by the modENCODE consortium was assembled. In addition to transcriptome profiling of worms at different developmental stages, the set included data from different mutant strains and worms exposed to a variety of pathogens. The raw sequencing reads for each experiment were treated in an identical fashion, with expression level reported as average depth of coverage per base per million reads (dcpm). The data, at a gene level in WS220, was converted to WS255 using WormBase Converter. Pearson correlations are calculated for the genes of interest within the dataset, the calculated correlations clustered as above and plotted using the function heatmap.2 within the R package ggplot.

### Fungal infection dataset

To define a test set of genes strongly induced by *D. coniospora* (“Dc_Up” genes), using published data (Table S7 in (Engelmann, et al., 2011)), we selected those with a >2.5- fold increase in infected versus control worms and with a minimum expression level of 0.1 dcpm in infected worms. The list of 145 genes was converted from WS170 to WS260 using WormBase Converter (Engelmann, et al., 2011), giving a list of 146 genes, including 2 that are non-coding.

## IMPLEMENTATION

### Constitution of reference gene sets for YAAT

In order to analyse a large dataset of genes derived from a genome-wide RNAi screen, we developed a functional class enrichment and clustering tool (Zugasti, et al., 2016). Given the interest generated by this tool, we decided to improve it and adapt it with the aim of providing a web-based resource, YAAT. The first improvement was to expand the range of underlying data. Our primary data source was Wormbase. From there, we extracted, for example, the genes associated with each of ca. 2000 phenotypic classes. A single gene can give rise to multiple transcripts, and hence to different protein isoforms. Some functional annotations are associated with a specific transcript, but we did not attempt to capture this level of detail in YAAT. Thus the reference object within YAAT is a gene (with unique Wormbase gene identifier; WBGene). Indeed, for annotations associated with proteins (e.g. for enzyme activities or derived from proteomic studies), the corresponding gene name was used. Thus for the sake of simplicity, here we use the term “gene” in an indiscriminate way when describing classes that were defined on the gene, transcript or protein level.

Wormbase has a regular release schedule, with updates available in principal every 2 months. Some data sources for YAAT are static. For example, the association of transcription factors and putative target genes established by the modENCODE consortium was generated using WS220 (Araya, et al., 2014) and although this large-scale study has been extended in the context of modERN (Kudron, et al., 2018), the modENCODE mapping of transcription factor targets has not been updated, despite subsequent changes in gene predictions. Other data sources, including KEGG (Tanabe and Kanehisa, 2012), are updated regularly. We previously developed a tool, Wormbase Converter, to deal specifically with this issue (Engelmann, et al., 2011). We used a script based on the Wormbase Converter algorithm to ensure that we had a homogeneous set of data, using a single reference release of Wormbase genes, irrespective of the data’s source.

Although we successfully performed one semi-automatic update of the YAAT database (from WS255 to WS260), with the passage of time, this became unfeasible for a number of reasons. One barrier to updating YAAT was changes in the location and availability of third party data. Common problems included out-dated URLs (e.g. the broken link to “a tab-delimited file of all public ‘hits’ (positive results) from DRSC screens” from https://fgr.hms.harvard.edu/utility-tools) and re-organisation of ftp data repositories. The use of database-specific gene identifiers (e.g. in KEGG) also presented a challenge as the inevitable occasional parsing errors required manual correction. Further, the release cycles of the different databases are not synchronised with Wormbase updates. For example, the Ensembl database is a variable number of releases behind Wormbase. This was particularly problematic for the establishment of phylogenetic profiles (see below) and could only be resolved by fastidious and very time-consuming cross-validation and error checking. Thus although YAAT was designed at the outset to be readily updated, we failed in that aim and have not yet been able to implement a robust pipeline for semi-automatic updating. Nevertheless, compared to many other tools, YAAT is based on relatively recent data and provides functions that they cannot. Two major sources of transcriptome data for YAAT were the pre-computed expression clusters from the *C. elegans* implementation of Serial Pattern of Expression Levels Locator (SPELL (Hibbs, et al., 2007)) and WormExp (Yang, et al., 2016). SPELL includes only published datasets, while the underlying data in WormExp comes from the many public transcriptome databases and includes unpublished data. These sources thus provided a partially redundant coverage of transcriptome data. We also expanded our pre-existing in-house functional class database (Engelmann, et al., 2011) by manually extracting data from transcriptome studies. Since this dataset only references gene identities, not expression values, it includes some studies for which the expression values were never made publicly available and that are therefore absent from SPELL and WormExp (Figure 1A). This transcriptome information was combined with functional and structural categories from different sources to constitute the YAAT database (Figure 1B).

**Figure 1.**
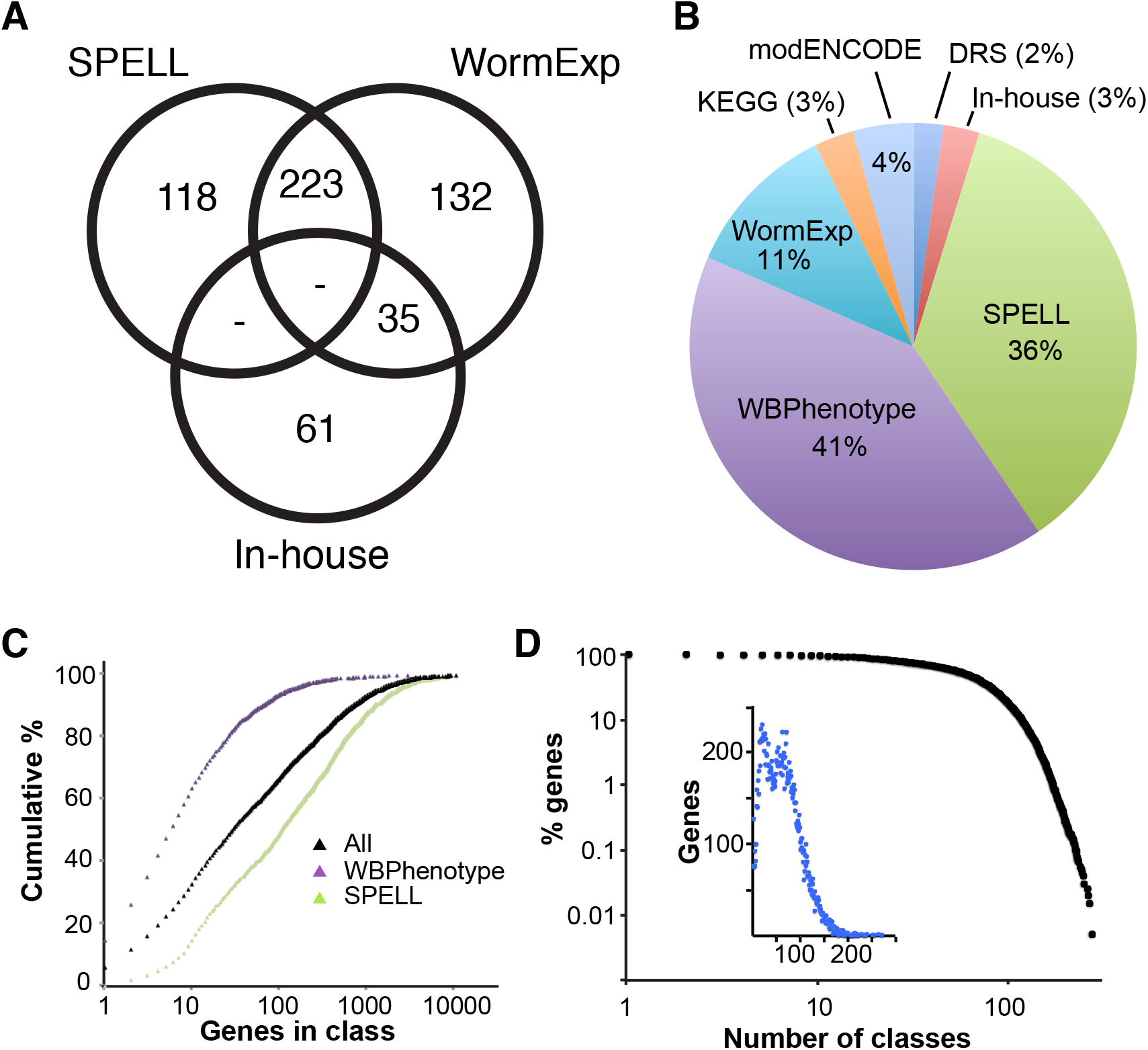
Sources and types of data in YAAT (WS255). **A**. Venn diagram showing transcriptome data coverage by SPELL, WormExp and our in-house collection. The figures indicate the number of articles from which data was extracted. **B**. Distribution of the major types of data in YAAT. DRS: phenotypes transposed from Drosophila RNAi screens. **C**. Distribution of class sizes, for all classes (black), and those from SPELL (orange) and Wormbase phenotypes (blue). **D**. Distribution of the number of classes associated with each gene. The inset graph is an alternative representation of the same data.

This set covers essentially all (>99%) protein-coding genes in C*. elegans*. The average class size was 272, with a median of 28 and a maximum of 10,705 genes (Figure 1C). Classes from different sources had markedly different size distributions. Thus the 1907 classes derived from Wormbase phenotypes were biased towards small classes (median 6, average 34), while those derived from SPELL (1662 classes) were biased towards large classes (median 104, average 484; Figure 1C).

Less than 130 genes (0.6%) were associated with a single category, in most cases (67/127 genes), the class “*C. elegans*-specific genes” (Zhou, et al., 2015). On the other hand, 95% of genes were associated with at least 10 classes, most frequently 21, with the median number of classes for each gene being 58 (Figure 1D). The complete dataset therefore has good gene coverage and the potential to provide insight into the functional relationships between the members of most gene lists.

### Gene sets analysis using YAAT

In its most basic implementation, a user inputs a list of genes of interest and sets analysis parameters. Users can choose between different methods to determine the statistical significance of each gene set enrichment; the default is by false discovery rate (FDR). Thresholds for FDR or P-values and for the maximum reference class size can be defined. In the context of enrichment analysis, including gene classes that contain as many as 50% of all protein-coding genes makes relatively little sense; we generally, and arbitrarily set this parameter to 1500 genes (i.e. ca. 7.5% of all protein-coding genes). Enriched functional classes are returned in a hyperlinked table that gives access to the underlying data. The results are also available as a downloadable text file and, as detailed further below, in different graphical formats.

### Analysis of the composition of the YAAT dataset reveals intrinsic biases

To characterise the dataset and the tool, we took the list of genes for each and every gene class and input this in turn into YAAT, to investigate the relationship between class sizes and number of enriched gene classes. We observed a positive correlatory trend between the number of genes in a class and the number of enriched functional classes for the entire set, and also for subsets corresponding to the Wormbase phenotype, WormExp or SPELL classes (Figure 2). There were, however, numerous classes that did not follow this trend. There were, for example, 4 large SPELL classes (of >5000 genes) associated with an unexpectedly low number of enriched categories (≤120, compared to a median of 416). It is not clear whether this reflects a technical artefact, but they come from a single study that transcriptionally profiled different cell types (Spencer, et al., 2011). At the other end of the scale, the class “Down <−1.5 by tunicamycin in N2”, with 126 genes returned more than 400 enriched classes (Table S1).

**Figure 2.**
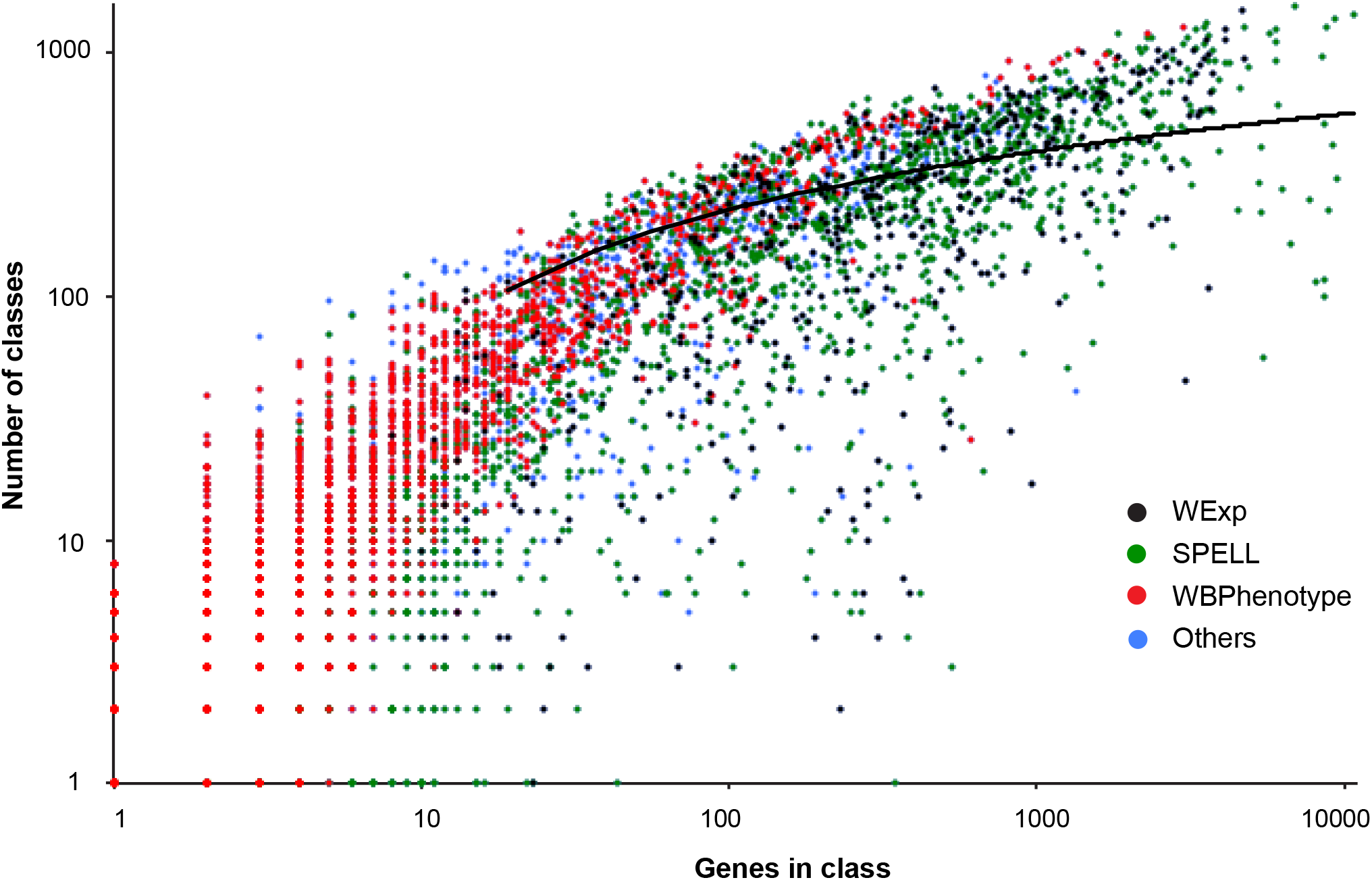
Distribution of the number of classes returned as a function of the size of the class used as a query, for classes derived from WormExp (black), SPELL (green), Wormbase phenotype (WBPhenotype; red) and others (blue). The line indicates the log fit using the entire set (WS255).

During the analysis, we noticed an apparent bias in the categories of classes returned for different gene lists. When the gene list used as a query was derived from a SPELL class, overall there appeared to be an over-representation of classes comprised of transcriptionally regulated genes (i.e. SPELL or WormExp). This was not seen with gene lists derived from WBPhenotype classes (Table S1). To probe this further, and to avoid the confounding effect of class size, we took the results for the 22 WBPhenotype classes constituted of 200-300 genes, and selected the 25 SPELL classes (only one per published study, chosen at random) of the same size that gave at least 300 enriched classes when run through YAAT (Figure 3A). Within these groups of gene lists, there was a striking difference in distribution of the enriched gene classes. While groups of genes defined by transcriptome studies (i.e. SPELL classes) were found to be enriched almost exclusively for other SPELL or WormExp classes, the groups defined on the basis of a phenotype were enriched for a far more diverse range of categories (Figure 3B), in proportions that approached their overall distribution (Figure 1B). Part of the reason for this result, as discussed further below, is that the respective gene “universes”, i.e. the non-redundant list of genes covered by all the classes of the various categories, are very different. The universe of SPELL or WormExp classes includes essentially all protein coding genes, while the other categories of classes only provide a partial coverage of the entire gene set. For example, most of the genes (57 ± 8 %, n = 13) in the SPELL-derived lists of 200-300 genes that returned the fewest (<5) WBPhenotype classes (Figure 3B), are not contained within the WBPhenotype universe of 8077 genes. The consistent bias that was observed is a caveat for the interpretation of this type of global enrichment analysis that draws its results from multiple types of gene annotations. In order to circumvent this potential issue, YAAT provides users with the possibility of restricting the enrichment analysis to defined categories of data.

**Figure 3.**
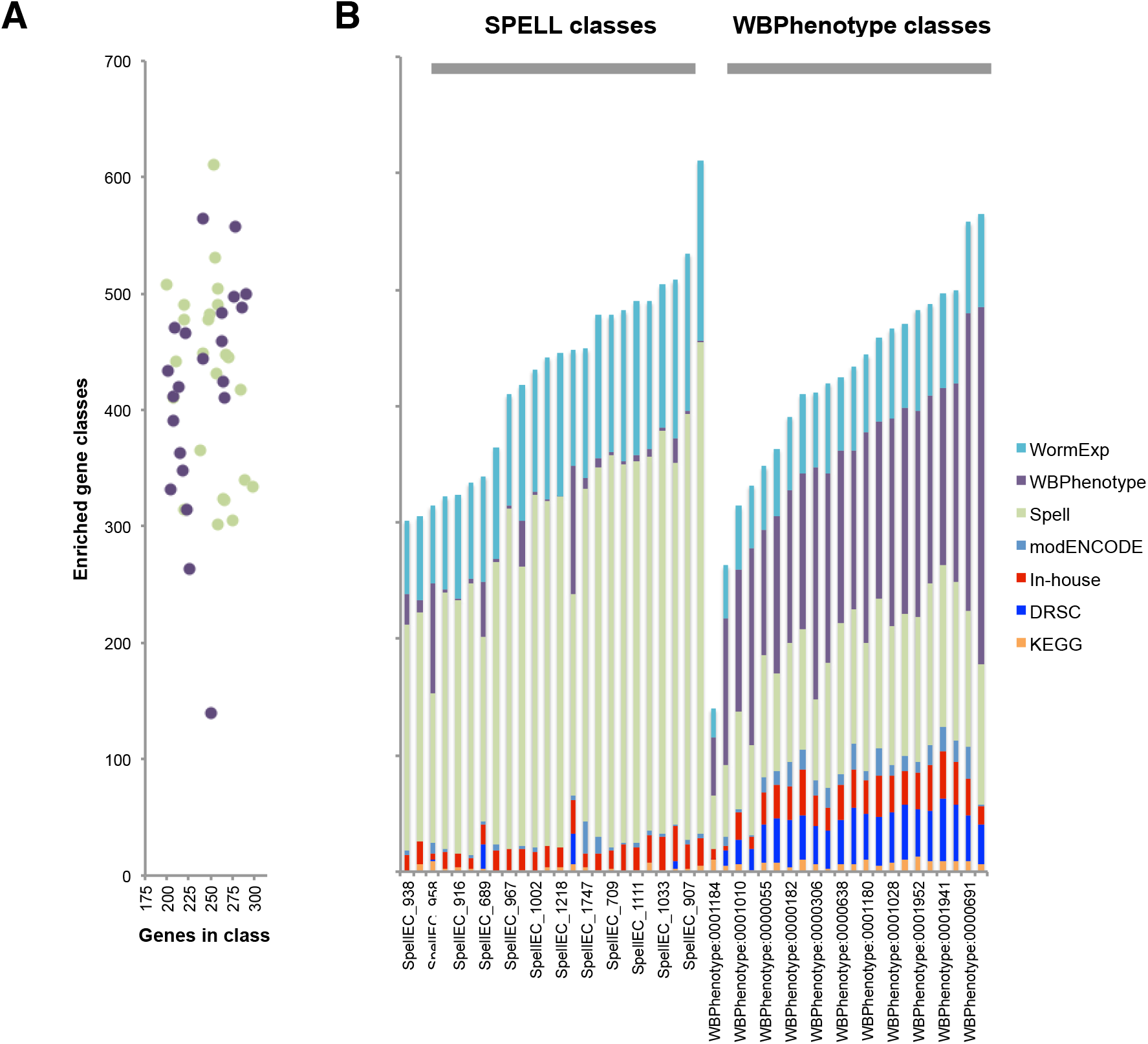
Sets of genes defined by transcriptome, but not phenotype, are preferentially enriched for certain gene classes. **A**. Characteristics of defined sets of genes from either the SPELL or WBPhenotype categories (see text for details). **B**. Graph showing the distribution of types of enriched gene classes for the same sets of genes. WS255 referenced data sets.

### Enriched class clustering to investigate functional connections

When a list of genes is entered, YAAT performs a standard enrichment analysis. To provide a graphical representation of the relationship between the members of the enriched classes, YAAT returns the results as a hierarchical cluster plot. Users can choose between 7 standard clustering methods, with Euclidean-distance based clustering being the default. The source data used for YAAT contains many related data sets. As a single example, from SPELL, there are 4 gene sets, “B.thuringiensis_0.5mix_upregulated_12h”, “B.thuringiensis_0.1mix_upregulated_12h”, “B.thuringiensis_0.5mix_upregulated_6h” and “B.thuringiensis_0.1mix_upregulated_6h” (Yang, et al., 2015) for which >70% of all the constituent genes are in more than one class, with 24% common to all 4. Such groups of related genes can strongly influence the clustering analysis and mask relationships that would otherwise be apparent between less-related classes. To counter this, YAAT offers the user the possibility to exclude classes that have too great an overlap from the analysis. Additionally, YAAT offers users the possibility to exclude any number of defined gene lists, which is useful, for example, when querying YAAT with part, or all, of a gene list that is in the database. Although hierarchical clustering is very powerful, and the utility of these analyses has been demonstrated previously (Kim, et al., 2016; Zugasti, et al., 2016), it can be biased because of the hierarchical nature of tree building. This can be avoided by calculating n-to-n distances between all classes and displaying the result as a network, where nodes represent enriched classes and edges represent the similarity between the different classes. YAAT therefore includes such a complementary network representation of these results, illustrating the degree of overlap of genes between different significantly enriched classes.

As an example, a set of 144 *C. elegans* protein-coding genes that are strongly up-regulated upon infection with the fungal pathogen *D. coniospora* (Dc_Up genes; see Methods), returns 39 highly enriched (p<10^−15^) gene classes. As expected from the global analysis of the dataset, there was a very strong bias of enriched classes; all were derived from transcriptome analyses (Table S2). In part this reflects the lack of representation of these genes in the other categories; only 34/144 were present in the WBPhenotype universe. Relaxing the threshold (P-value < 0.01) revealed the common functional annotation, “pathogen susceptibility increased” for 5 of these genes (*fmo-2; thn-1; thn-2; ilys-3; Y65B4BR.1*), since their individual or compound knockdown was associated with loss of resistance to one or more pathogens (Evans, et al., 2008; Luhachack, et al., 2012; O’Rourke, et al., 2006; Sahu, et al., 2012; Shapira, et al., 2006). They are likely to form part of a general but relatively unexplored pathogen defence strategy that is the subject of current investigation (Dasgupta, et al., 2020; Naim, et al., 2021). This illustrates one way in which YAAT can be used to generate new knowledge at the gene level.

YAAT provides a link to the sets of genes shared between the query list and each enriched class, allowing iterative rounds of analysis, to explore in more depth the relationships between classes. Since the results are available as a downloadable text file, users can represent the results graphically, such as with volcano plots. To provide a graphical representation of the relationship between the members of the enriched classes, YAAT returns the results as a hierarchical cluster plot. With the Dc_Up set of query genes, there were several distinct clusters. Within them, a number of classes clustered as expected. For example, the genes regulated upon *D. coniospora* infection and in the osmotic stress-related mutants *osm-7* and *osm-11* overlap significantly (Rohlfing, et al., 2010) and so co-clustered (Figure 4A; Table S2).

**Figure 4.**
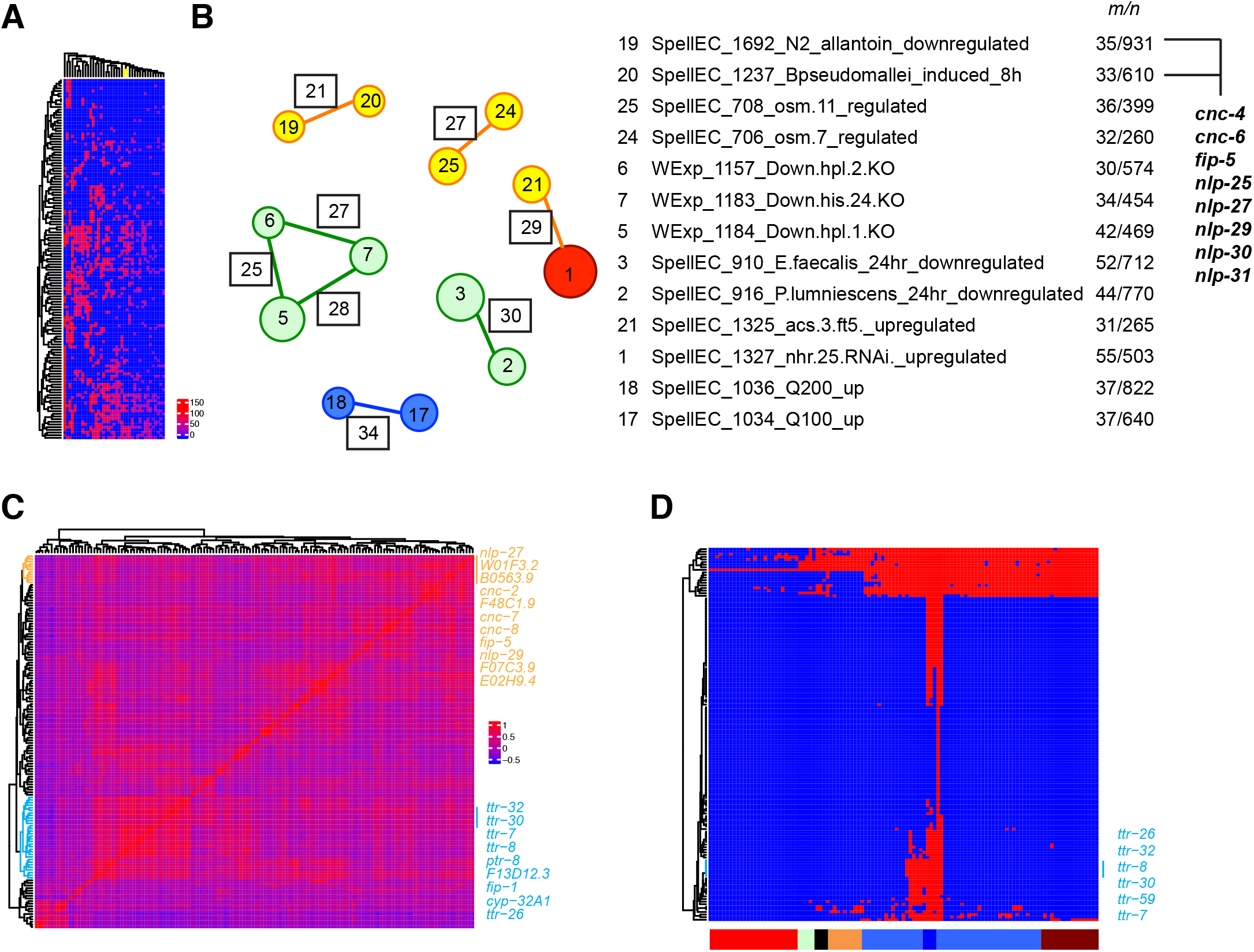
**A**. Clustering of enriched classes for 144 *D. coniospora* up-regulated genes. Each row is a gene, each column is a class; the labels are not shown here. The positions on the dendogram of the classes “SpellEC_706_osm.7_regulated” and “SpellEC_708_osm.11_regulated” is indicated in yellow. The colour scale indicates the number of genes in the overlap between the input gene list and each enriched class. **B**. Network of enriched gene classes. The colour of each node (gene class) corresponds to a position in the hierarchical cluster, its size indicates the significance (p-value) of the overlap between the class and the query list of genes (larger diameter, higher significance). In YAAT, the gene class labels are reported directly in the network. Here, they are replaced by a number in each node (in their order, from left to right, in the graph in **A**, with the correspondence table on the right), for the sake of legibility. The boxes indicate the number of genes shared between each pair of gene classes and the query list. The correspondence table gives the gene class, with *m*/*n*, *n* the total number of genes of that class and *m* the number that overlap with the query set. The righthand list of genes highlights antimicrobial peptide genes common to the overlaps of classes #19 and #20. **C**. Clustering of 144 *D. coniospora* up-regulated genes on the basis of their patterns of co-expression. Two subclusters are highlighted, with the positions of the indicated genes marked by the vertical lines on the right. The colour scale indicates the degree of co-expression between gene pairs. **D**. Clustering of 144 *D. coniospora* up-regulated genes on the basis of their patterns of conservation (red indicates the presence of an orthologue) across 113 species (coloured bar at bottom), ranging from Archaea and bacteria (red; left) to chordates, including human (brown; right). Among the invertebrates (blue), the 5 *Caenorhabditis* species are indicated by the darker blue box. One gene subcluster is highlighted, with the positions of the indicated genes marked by the vertical line on the right. The query parameters and full results for all panels are in Table S2.

When the similarity between the enriched classes is calculated and then represented in the form of a class linkage network, the connections between a more select group of classes becomes evident (Figure 4B). In this case, the analysis highlights 3 known connections with the anti-fungal innate immunity, again, the overlap with the response to osmotic stress (Dodd, et al., 2018; Pujol, et al., 2008; Rohlfing, et al., 2010; Zugasti, et al., 2016); the reciprocal relationship with the response to bacterial pathogens that colonize the intestine (Engelmann, et al., 2011); and the link to *acs-3*/*nhr-25* signalling (Ward, et al., 2014). These last 2 categories were not in the same sub-cluster in the dendrogram (Figure 4A, Table S2), illustrating the utility of this complementary analysis. There was also an overlap with lists corresponding to genes induced upon exposure to 2 concentrations of the bioactive polyphenol quercetin acid (Pietsch, et al., 2012), in part explicable by the >90% overlap between these 2 latter classes. Perhaps more interestingly, this representation revealed an overlap between the *Drechmeria*-induced genes and genes up-regulated by *Burkholderia pseudomallei* infection (Lee, et al., 2013) but repressed by allantoin (Calvert, et al., 2016). The commonly regulated genes include a hallmark of epidermal antifungal immunity, 8 antimicrobial peptide genes (Figure 4B). *B. pseudomallei* degrades ELT-2, an intestinal GATA transcription factor, and so switches off defence gene expression in the gut (Lee, et al., 2013). How this would switch on immune gene expression in the epidermis is not known. Allantoin extends *C. elegans* lifespan in a DAF-16-independent manner through a mechanism that is not fully understood (Calvert, et al., 2016). The potential for allantoin to modulate innate immunity merits further investigation. While the results with this test dataset illustrate the confounding effect of including highly-related gene lists in the database, more importantly they suggest that YAAT has the potential to identify functional connections linking different regulatory mechanisms.

### Establishing functional connections using co-expression analysis

Genes that function together often are expressed together. Therefore, as well as calculating and returning measures of gene class enrichment, YAAT generates, from a set of ModENCODE RNAseq data (Agarwal, et al., 2010), information about gene co-expression. Taking the same set of 144 Dc_Up genes, a number of separate clusters can be resolved, including one that includes several antimicrobial peptides genes (Figure 4C). In addition to the annotated *cnc*, *fip* and *nlp* genes (Pujol, et al., 2008), this clustering draws attention not only to *F48C1.9*, recently demonstrated to be up-regulated specifically in the epidermis upon *D. coniospora* infection (Omi and Pujol, 2019) but also to *B0563.9* predicted to encode a secreted peptide of around 50 amino acids structural unrelated to previously characterised antimicrobial peptides, and *F07C3.9*, predicted to encode a 53 amino acid peptide, without a signal peptide, with homologues only in some *Caenorhabditis* species. Both of these merit further study. Another clearly discernable cluster contained 5 *ttr* (TransThyretin-Related family domain) genes (Figure 4C). Several members of this large family of 59 genes have been implicated in *C. elegans* host defence (e.g. (Simonsen, et al., 2011; Treitz, et al., 2015)), but their role in the interaction with *D. coniospora* has not yet been investigated. These results clearly illustrate how YAAT can help prioritise genes for in-depth study.

### Establishing functional connections using phylogenetic analysis

Genes that function together are also often observed to co-evolve (Pellegrini, et al., 1999). To provide a comparative measure of gene conservation, YAAT displays the average level of conservation across a broad range of species for the entire query list. It also includes in the table of results a comparison between the average level of conservation for the entire list of genes in an enriched class and for the genes that are shared between that list and the genes input as a query. The figures for overall conservation for the members of the different classes varied widely. There was a strong bias towards conservation among the “phenotype” classes but not among those derived from the SPELL expression studies (Table S3). Not surprisingly, among the larger classes (>200 genes), the lowest score was associated with the class of lineage-specific genes (Zhou, et al., 2015). It was notable that 6 of 30 classes with the least conserved genes were lists of infection-regulated genes, presumably reflecting the lineage-specific adaptation to selection pressure from natural pathogens. At the other end of the scale, with high scores, apart from the classes transposed from *Drosophila*, which were by their very nature conserved, those related to fundamental cellular processes, including KEGG gene classes, were prominent. The observed skew in the conservation of genes found in the different types of functional classes could lead to another type of bias when analysing datasets.

In addition to giving a global measure of conservation, to provide insight into the pattern of conservation of the members of a gene list, YAAT performs phylogenetic profiling. By default, the results of a broad analysis against 113 diverse species are presented. Given the increasing interest in translating research from *C. elegans* to parasitic worms (e.g. (Keiser, 2015)), users can opt for a more targeted one, using data from 127 nematode and helminth species. With the list of 144 Dc_Up, most genes were either found only in nematode species or just in *C. elegans* (Figure 4D). The 5 *ttr* genes mentioned above were found in the same sub-branch of the phylogenetic tree, increasing the likelihood that they function together in an unexplored aspect of innate defence.

### Simultaneous clustering of class enrichment, co-expression and phylogenetic data

Genes that have the same pattern of class membership, co-expression and conservation may very well function together. To facilitate the identification of such groups of genes, YAAT uses a powerful new form of representation. For each set of class enrichment, co-expression and phylogenetic data, the order of genes in the clustering is enforced for the results of the other two sets. To demonstrate the utility of the approach, we queried YAAT with a combined list of genes corresponding to 2 large protein complexes with no common members (Table S4). Clustering by class enrichment resulted in a very good separation of the 2 lists. When this clustering was enforced on the co-expression data, clear groups were discernable (Figure 5A). Similarly, clustering on the basis of co-expression separated the genes from the 2 complexes in a very satisfactory manner, while projection of this clustering on the class enrichment data also resulted in clear groups of genes (Figure 5B). With this data set, clustering on the basis of evolutionary conservation separated the 2 gene sets relatively poorly, presumably as a consequence of the fact that almost all the genes, regardless of the set, exhibit the same pattern of conservation. There was one marked exception, a group of genes specific to only one complex with a unique pattern of conservation (Figure 5C) that could represent the ancestral core complex.

**Figure 5.**
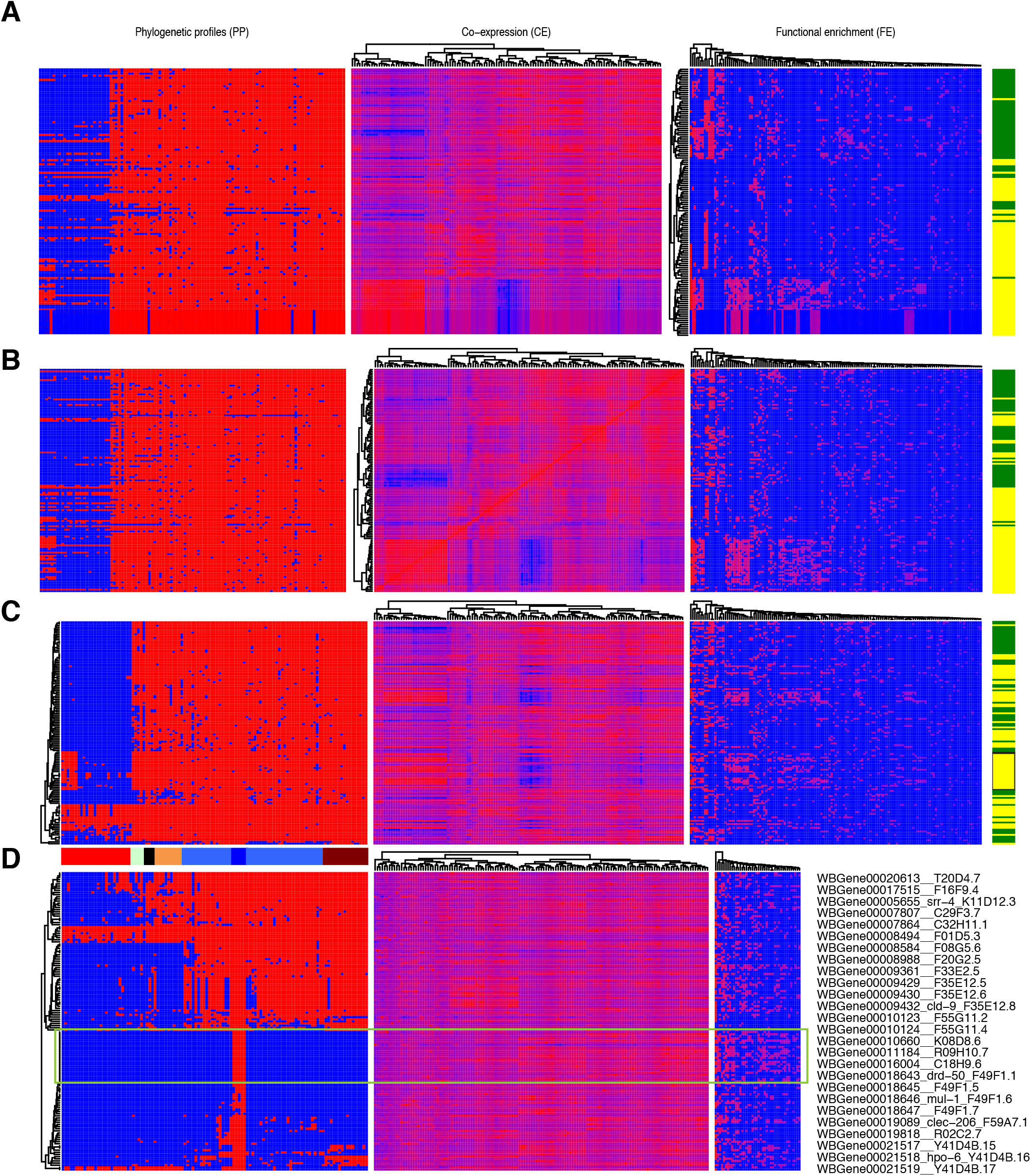
**A**. Clustering of enriched classes for genes from 2 independent protein complexes, indicated by the green and yellow bars on the right. Each row is a gene, each column is a class; the labels are not shown here. The gene order obtained from the class clustering (right panel) is imposed on the co-expression (middle panel) and conservation (left panel) data. Clustering of the same genes on the basis of co-expression (**B**) or conservation (**C**; one group of genes is highlighted by the black box on the right; species colour code as in Figure 4), with in each case the respective gene order being imposed on the 2 other panels. **D**. Clustering of genes induced by exposure to RPW-24 on the basis of conservation (left panel), with this gene order being imposed on the 2 other panels. The identity of the boxed group of genes is shown on the right. The query parameters, the full species colour code and full results for all panels are in Tables S4 and S5.

As another example, we took a published list of genes determined to be induced by the immunostimulatory xenobiotic RPW-24 in an *nhr-86*-dependent manner (Peterson, et al., 2019). In this case, there was a broader phylogenetic distribution. Looking at the clustering based on evolutionary conservation, one can see a coherent group of genes that were specific to the 5 *Caenorhabditis* species that also gave a clear cluster of 26 genes across the other 2 graphs (Figure 5D). Despite representing less than 1/5 of the input list, these genes were the majority of those found in 2 classes, “N2_UV_upregulated” and “PA14_vs_OP50_upregulated_8hr” (Table S5). As could be expected, at the gene level, 8 had annotations linked to the response to a range of bacterial pathogens, and 4 to abiotic stress. Interestingly, several correspond to proteins of the same family, in close proximity to each other in the genome (Table S5). These are all indications for recently evolved genes potentially in cellular defence, and again illustrate how YAAT can help identify groups of genes for in-depth functional study.

## DISCUSSION

Various stand-alone or web-based functional enrichment analysis tools have been developed over the last 20 years. The results of our tests with YAAT indicate that enrichment analyses should be based on comparisons with the broadest possible collection of functional classes to avoid the biases inherent to each type of gene set. Indeed, by collecting transcriptomic, functional and phenotypic data from the literature, Wormbase, KEGG and other resources, we amassed more than 10,000 gene classes, covering essentially all protein coding genes in Wormbase. Since data evolves, from the outset, we included a pipeline to update data from the respective resources automatically. For the static data such as modENCODE assigned transcription factor targets, the most accurate updating would involve reassigning genes to ChIP-seq peaks on the basis of revised gene structure predictions. Even for RNA seq data, each revision of gene predictions should be accompanied by a re-calling of alignments and read counts. Such practices are today unfeasible. Indeed, even our intermediate strategy, tracking changes in gene identity (and associated annotations) over successive Wormbase releases using an established methodology (Engelmann, et al., 2011) proved unworkable after just one update cycle. We are well aware of the material and theoretical difficulties in maintaining any biological database, but have no concrete solutions to the multiple problems we encountered trying to keep YAAT up to date. Regarding the analyses themselves, rigorous statistical enrichment analysis requires knowledge of the gene “universe” sampled when a class was defined. It matters whether a class was defined after sampling all genes or only a subset. For example, in our previous transcriptome studies to determine the response of *C. elegans* to infection, we have used cDNA (Mallo, et al., 2002), long oligonucleotide (Wong, et al., 2007) and tiling microarrays (Engelmann, et al., 2011). These theoretically cover between 35% and close to 100% of the predicted protein coding genes. In reality, because of the sensitivity of the techniques, these figures are upper limits. Similarly, for analyses by RNA-seq, in principal all genes are assayed, but in fact the number is determined by the depth of sequencing (Tarazona, et al., 2011), and in some studies may be restricted to a particular category. For example, a recent study looked specifically for non-coding transcripts regulated by CEP-1 (Xu, et al., 2014). All of the genes identified in this study that are associated with a Wormbase SPELL class included in YAAT have both protein-coding and non-coding transcripts. Currently <550 such genes are predicted, and this would be the relevant universe for these classes, not the >25,000 non-protein-coding genes. For other types of class, determining the universe is problematic, be it because of the high rate of false negatives in RNAi screens, or lack of relevant information (i.e. degree of saturation) for genetic screens. Indeed, it is very rare for the sampled universe to be reported, and even when it is, this information is generally not available in a machine-readable standardised format. In common with most current functional enrichment analysis tools, in YAAT we assumed that the universe for each set is the entire complement of protein coding genes. This will introduce inevitable inaccuracy in the calculation of the statistical significance of any given enrichment, especially if the universe of a class of interest is small.

YAAT goes beyond the functionality of standard enrichment analysis since it provides class clustering, allowing the inference of functional links between groups of genes. Further, we have included the option to perform co-expression and phylogenetic profiling, complementary methods for establishing groups of functionally-related genes. YAAT was originally provided as a web-based service on an academic web server. Following alteration to the host site’s security protocols, external access was lost. Despite our best efforts over many months, we have not succeeded in reinstalling YAAT elsewhere, in part because of the complexity of the underlying programming environment with the attendant multiplicity of dependencies. Unfortunately, YAAT was conceived before the widespread implementation of containers like Singularity that are specifically designed to circumvent such constraints. We will continue to try to restore and online version of YAAT and implement methods to allow it to be updated. Even if we do not succeed, we hope that our work with YAAT will serve as a model for the type of integrated analysis tool that could be useful for any large-scale exploration of biological function as well as illustrating the various pitfalls that can plague meaningful interpretation of gene lists.

## Acknowledgements

Supported by institutional grants from the Institut national de la santé et de la recherche médicale, Centre national de la recherche scientifique and Aix-Marseille University to the CIML, and the Agence Nationale de la Recherche program grants (ANR-16-CE15-0001-01, ANR-11-LABX-0054 (Labex INFORM) and ANR-11-IDEX-0001-02 (A*MIDEX)). We thank Damien Courtine and Lionel Spinelli for their input.

